# Disinhibition as a canonical neural mechanism for flexible behavior

**DOI:** 10.1101/334797

**Authors:** Dominic Standage, Martin Paré, Gunnar Blohm

**Affiliations:** Centre for Neuroscience Studies; 18 Stuart Street; Kingston, ON, K7L 3N6, Canada

**Keywords:** decision making, working memory, network

## Abstract

Flexibility is a hallmark of human and animal behavior, but the context-dependent neural computations that generate flexible behavior are poorly understood. Here, we use a biophysically-based cortical network model to explore the hypothesis that vasoactive intestinal polypeptide (VIP) expressing inhibitory interneurons control local circuit dynamics by targeting other classes of inhibitory interneuron, supporting context-dependent computations. Depending on the strength of this disinhibition (simulating VIP activity), network dynamics support multiple-item working memory (WM, strong disinhibition) or decision making (DM, weak disinhibition). Within these regimes, disinhibition controls WM capacity and speedaccuracy-trade-off in choice behavior. Our findings suggest that long-range trans-cortical VIP-mediated disinhibition is a canonical neural mechanism for the top-down control of flexible behavior.

## Introduction

Recent data (Pi et al., 2013; Coxon, Peat, & Byblow, 2014) provide evidence for a highly conserved cortical micro-circuit motif specializing in disinhibition. Within this motif, VIP interneurons target other inhibitory interneurons (parvalbumin and somatostatin expressing cells) that in turn synapse onto pyramidal cells. The functional role and significance of this form of disinhibition is unknown. but it has been hypothesized that disinhibitory control gates information transmission (Wang & Yang, 2018) and tunes cortical processing capabilities (Coxon et al., 2014); and that disregulation of disinhibition might underlie abnormal attention (Pezze, McGarrity, Mason, Fone, & Bast, 2014) and sensory processing (Nunes & Kuner, 2015). Because VIP interneurons are directly targeted by distal brain areas, as well as by neuromodulators, they are strong candidates for implementing flexible control over local circuit behavior.

It is widely believed that several association cortical areas (e.g. dorsolateral prefrontal cortex and the lateral intraparietal area) play an important role in DM and WM, but these two cognitive processes make conflicting demands of neural circuitry. Competition between neural populations encoding choice alternatives is crucial to DM (Wong, Huk, Shadlen, & Wang, 2007; Standage & Paré, 2011), but competition between populations encoding memoranda entails forgetting. Thus, a mechanism is required to support both processing regimes. To the best of our knowledge, no such mechanism has been proposed. Here, we hypothesize that VIP disinhibitory control induces rapid transitions between network regimes. To instantiate this hypothesis, we simulated disinhibitory control in a network model.

## Methods

Our local-circuit model is a network of pyramidal neurons and inhibitory interneurons, connected by AMPA, NMDA and GABA receptor conductance synapses (Fig. 1). We simulated two tasks: a luminance-contrast discrimination task and a WM task. On the former (decision) task, the network distinguished a higher-rate stimulus (the target) from a lower-rate stimulus or stimuli (the distractors). On the memory task, the network accurately retained as many stimuli as possible over a delay interval. Cognitive control by disinhibition was simulated by scaling a non-selective inhibitory conductance onto interneurons, mimicking the effect of VIP interneurons on both tasks.

**Figure 1:**
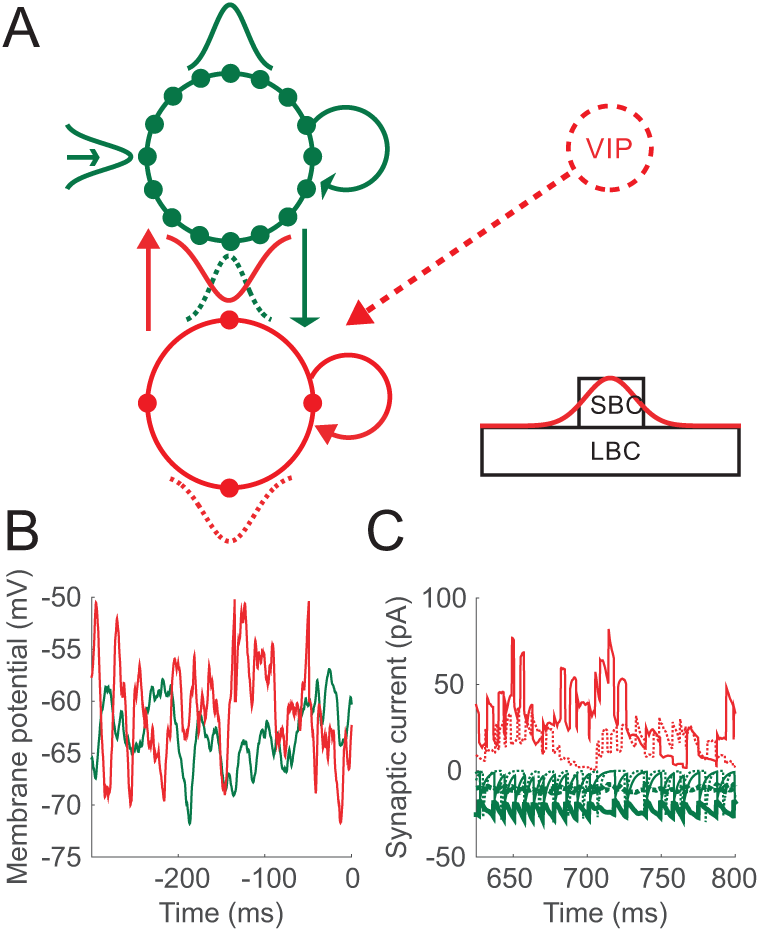
(A) Local-circuit model. Periodical pyramidal neurons (solid green) and inhibitory interneurons (red). Arrows depict connectivity (Gaussian) and RF (Gaussian with arrow). GABAR (red arrows), AMPAR-only (open green arrows), and synapses with AMPARs and NMDARs (wide green arrows). Inhibition from small and large basket cells (SBC and LBC). (B) Membrane potential of a pyramidal neuron and an interneuron at rest. (C) Synaptic currents (red: GABAR; thin green: AMPAR; thick green: NMDAR) onto a pyramidal neuron (solid) and an interneuron (dotted).

The local circuit model is a fully connected network of leaky integrate-and-fire neurons (*N*^*p*^ = 800 pyramidal, *N*^*i*^ = *N*^*p*^/4 fast-spiking inhibitory interneurons). Each model neuron is described by

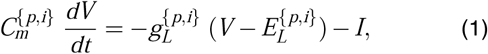

where *C*_*m*_ is the membrane capacitance, *g*_*L*_ is the leakage conductance, *V* is the membrane potential, *E*_*L*_ is the equilibrium potential, and *I* is the total input current. When *V* reaches a threshold ϑ_*v*_, it is reset to *V*_*res*_, after which it is unresponsive to its input for an absolute refractory period of τ_*ref*_. Here and below, superscripts *p* and *i* refer to pyramidal neurons and interneurons respectively. The input current at each neuron is

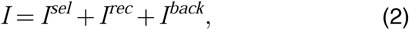

where *I*^*sel*^ is stimulus-selective synaptic current (0 for interneurons), *I*^*rec*^ is recurrent (intrinsic) synaptic current and *I*^*back*^ is background current. *I*^*sel*^ and *I*^*rec*^ are synaptic currents, and *I*^*back*^ is injected current. Synaptic currents driven by pyramidal neuron spiking are mediated by simulated AMPA receptor (AMPAR) and/or NMDA receptor (NMDAR) conductances, and synaptic currents driven by interneuron spiking are mediated by simulated GABA receptor (GABAR) conductances. Synaptic activation (proportion open channels) is

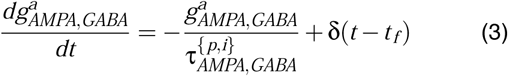

where τ _*AMPA*_ and τ_*GABA*_ are the time constants of AMPAR and GABAR deactivation respectively, δis the Dirac delta function, *t*_*f*_ is the time of firing of a pre-synaptic neuron and superscript *a* indicates that synapses are activated. NMDAR activation has a slower rise and decay

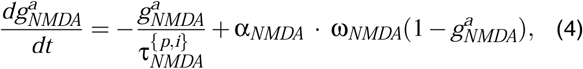

where τ _*NMDA*_ is the time constant of receptor deactivation and α _*NMDA*_ controls the saturation of NMDAR channels at high pre-synaptic spike frequencies. The slower opening of NMDAR channels is captured by

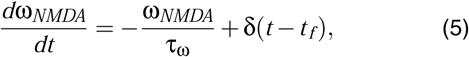

where τ_ω_ determines the rate of channel opening. Intrinsic (recurrent, local feedback) synaptic current to neuron *j* is

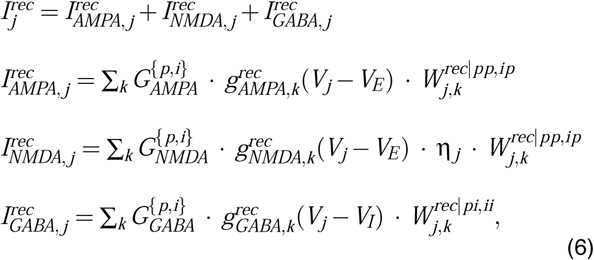

Matrices *W*^*rec*|*pp*,*ip*^ and *W*^*rec*|*pi*,*ii*^ scale conductance strength or *weight* according to network connectivity. Gaussian distance-dependent connectivity depends on the class of neuron receiving and projecting spiking activity, where superscripts *pp*, *ip*, *pi* and *ii* denote connections to pyramidal neurons from pyramidal neurons, to interneurons from pyramidal neurons, to pyramidal neurons from interneurons, and to interneurons from interneurons respectively. The weight 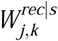 to neuron *j* from neuron *k* is given by

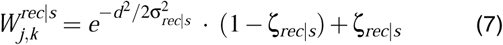

where *d* is the distance between preand post-synaptic neurons. Parameter σ_*rec*|*s*_ determines the spatial extent of connectivity and parameter ζ_*rec*|*s*_ allows the inclusion of a baseline weight, with the function normalized to a maximum of 1 (0 ≤ ζ_*rec*|*s*_ < 1).

Cortical background activity was simulated by the point-conductance model (Destexhe, Rudolph, Fellous, & Sejnowski, 2001):

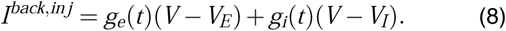

The excitatory and inhibitory conductances *g*_*e*_(*t*) and *g*_*i*_(*t*) are updated at each timestep Δ*t* according to

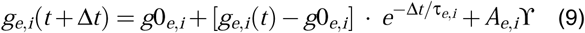

where *g*0_*e*_ and *g*0_*i*_ are average conductances, τ_*e*_ and τ_*i*_ are time constants, and ϒ is normally distributed random noise with 0 mean and unit standard deviation. Amplitude coefficients *A*_*e*_ and *A*_*i*_ are defined by

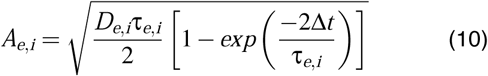

where 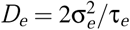 and 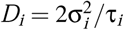 are noise ‘diffusion’ coefficients (see Destexhe et al. (2001)).

For inhibitory interneurons, the background conductance *g*_*i*_(*t*) was further used to simulate the activity of a population of VIPs. To this end, the average inhibitory conductance *g*0_*i*_ was scaled by control parameter *G*_*ii*_ for all interneurons (0.6 ≤ *G*_*ii*_ ≤ 4), implementing the hypothesized disinhibitory signal (Figures 1A).

We simulated the target stimuli in both tasks by providing independent, homogeneous Poisson spike trains to all pyramidal neurons *j* in the network, where spike rates were drawn from a normal distribution with mean *µ*_*sel*_ corresponding to the center of a Gaussian response field (RF) defined by 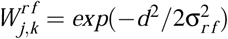. Constant *d* is given above for recurrent synaptic structure *W*^*rec*|*pp*^, σ_*r f*_ determines the width of the RF and subscript *k* indexes the neuron at the RF center. Spike response adaptation by upstream visually responsive neurons was modeled by a step-and-decay function. The stimuli were mediated by AMPARs only, so for all pyramidal neurons *j* in the PPC network,

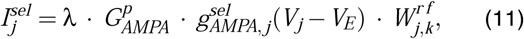

where constant λ scales extrinsic synaptic conductance. All simulations were run with the standard implementation of Euler’s forward method and a timestep of Δ*t* = 0.25ms.

## Results

Changing the strength of disinhibition (*G*_*ii*_) produced network behavior supporting DM (left column in Fig. 2) and WM (middle and right columns).

**Figure 2:**
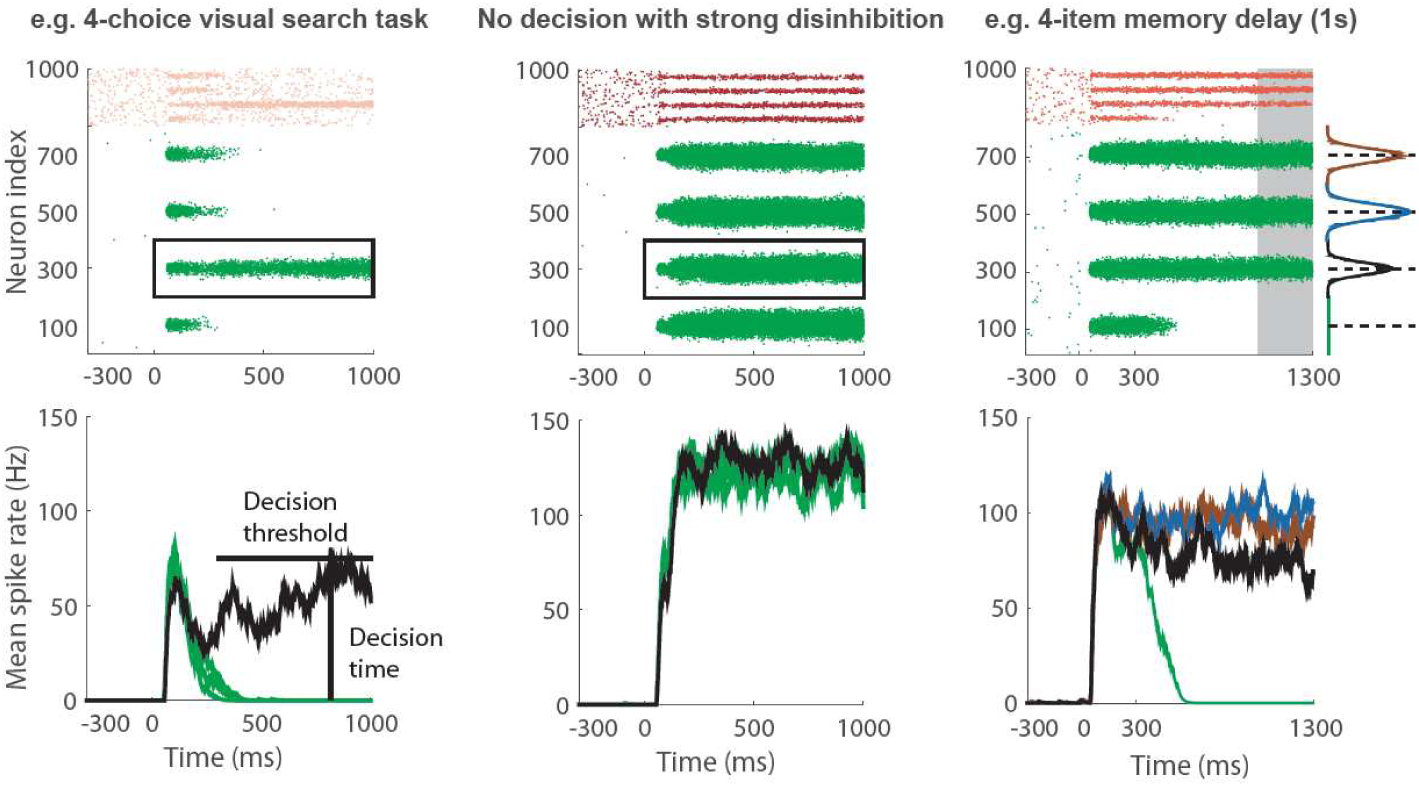
Network simulations with different strengths of disinhibitory control *G*_*ii*_.

### Disinhibition controls DM

Low values of *G*_*ii*_ support decision-making behavior. Outside a certain range, feedback inhibition was too weak or too strong to support competitive dynamics. We refer to the range of *G*_*ii*_ that supported DM as 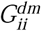.

Consistent with electrophysiological recordings from monkey PPC (Churchland, Kiani, & Shadlen, 2008), the mean firing rate of the initial featureless response decreased with an increase in the number of stimuli (*n*) for all 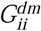 and all task difficulties; the slope of decision-selective activity was lower with higher task difficulty in the lead-up to decision time for all 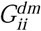 and all *n*; and the slope of this activity increased with *n* for a given task difficulty for all 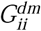 (data not shown). Consistent with behaviour, the model made fast, accurate decisions on easy trials; and slower, less accurate decisions as task difficulty was increased for all 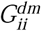, *i.e.* the model showed typical psychometric and chronometric curves (Fig. 3).

**Figure 3:**
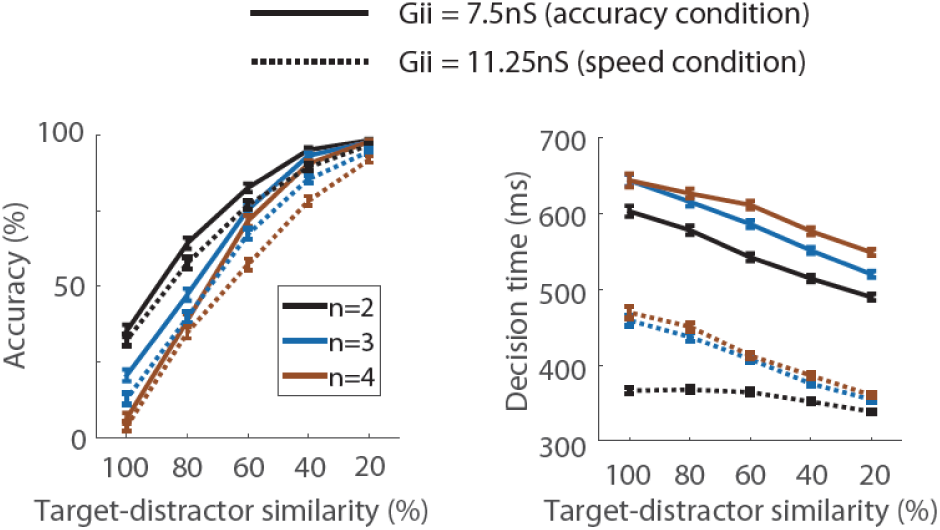
Disinhibition controls speed-accuracy trade-off for different number of choices *n*.

Not only did disinhibition support DM in a manner consistent with neural and behavioural data, but it further controlled the speed-accuracy trade-off (SAT) on the decision task. For a given task difficulty, decisions were faster and less accurate with higher *G*_*ii*_ (Fig. 3). Thus, disinhibition not only offers a plausible mechanism by which generic cortical circuitry can be modulated to support DM, but further offers a mechanism for controlling decision processing according to task conditions. Such flexible cognitive control is fundamental to choice behaviour [see Standage, Wang, and Blohm (2014)].

### Disinhibition controls WM capacity

The network supported WM storage for higher values of 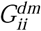. Simulated neural activity on the 1-item memory task qualitatively reproduced single-cell recordings from monkey PPC, showing a rapid-onset response during the stimulus interval, the rate of which exceeded the steady-state rate during the delay interval (Paré & Wurtz, 1997) (data not shown).

To measure WM performance for each value of 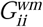, we defined capacity *K*(*n*) as the mean number of accurately retained items for each *n*-item memory task, and we defined peak capacity as the maximum value of *K*(*n*). We refer to the value of *n* corresponding to peak capacity as 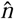 and we refer to a decrease in capacity for 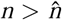 as WM overload (Matsuyoshi, Osaka, & Osaka, 2014). With moderately strong disinhibition, peak capacity was consistent with that of monkeys [2 ± 1 (Heyselaar, Johnston, & Paré, 2011)] and humans [4 ± 1 (Luck & Vogel, 1997; Cowan, 2001)] (Fig. 4B). Peak capacity increased very slightly for higher *G*_*ii*_, but WM overload became unrealistically pronounced, with capacity dropping from *K*(4) ≈ 4 to *K*(5) ≈ 0.5. With very strong disinhibition, peak capacity decreased and overload remained pronounced, providing a functional ‘ceiling’ on *G*_*ii*_, *i.e.* there was no benefit to stronger disinhibition (Fig. 4C).

**Figure 4:**
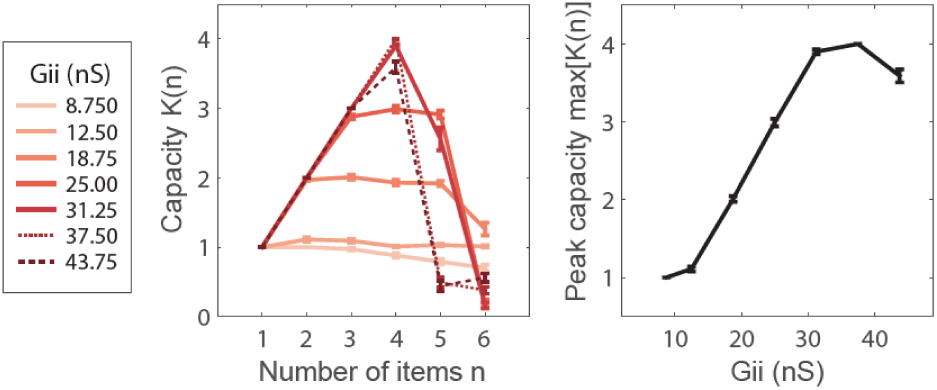
WM performance. Capacity (left) depends on number of items *n*. Peak capacity (right) as a function of *G*_*ii*_.

## Conclusion

In this proof-of-principle study, we showed that VIP-mediated disinhibition is a potential canonical mechanism for flexible cognitive control. Furthermore, disinhibition effectively changed the RF of stimulus-selective pyramidal cells (width and strength, data not shown), consistent with the effect of top-down attentional control (Martinez-Trujillo & Treue, 2004).

## Acknowledgments

Supported by NSERC and CFI (Canada).

